# Fingerprinting of skin cells by live cell Raman spectroscopy reveals melanoma cell heterogeneity and cell-type specific responses to UVR

**DOI:** 10.1101/2021.11.12.468396

**Authors:** Emma L. Wilkinson, Lorna Ashton, Jemma G. Kerns, Sarah L. Allinson, Richard L. Mort

## Abstract

Raman spectroscopy is an emerging dermatological technique with the potential to discriminate biochemically between cell types in a label free and non-invasive manner. Here we use live single cell Raman spectroscopy and principal component analysis (PCA) to fingerprint mouse melanoblasts, melanocytes, keratinocytes and melanoma cells. We show the differences in their spectra are attributable to biomarkers in the melanin biosynthesis pathway and that melanoma cells are a heterogeneous population that sit on a trajectory between undifferentiated melanoblasts and differentiated melanocytes. We demonstrate the utility of Raman spectroscopy as a highly sensitive tool to probe the melanin biosynthesis pathway and its immediate response to UV irradiation revealing previously undescribed opposing responses to UVA and UVB irradiation in melanocytes. Finally, we identify melanocyte specific accumulation of β-carotene correlated with a stabilisation of the UVR response in lipids and proteins consistent with a β-carotene mediated photoprotective mechanism. In summary our data show that Raman spectroscopy can be used to determine the differentiation status of cells of the melanocyte lineage and describe the immediate and temporal biochemical changes associated with UV exposure which differ depending on cell type, differentiation status and competence to synthesise melanin.

## INTRODUCTION

Melanocytes are the pigment-producing cells found in the skin, hair and eyes. Their embryonic precursors are melanoblasts derived from a transient tissue known as the neural crest (Mort et al. 2015). Melanocytes manufacture melanin from tyrosine and phenylalanine using the enzymes phenyalanine hydroxylase (PAH), tyrosinase (TYR), dopachrome tautomerase (DCT) and tyrosinase related protein 1 and 2 (TYRP1 and TYRP2) (Schallreuter et al. 1994). They use their dendrites to export melanin containing melanosomes to keratinocytes which arrange them above their nuclei to protect from ultraviolet radiation (UVR) induced DNA damage (Kobayashi et al. 1998; Maddodi et al. 2012). Melanoma, the cancer of melanocytes, is the deadliest form of skin cancer with a rising worldwide incidence (Erdei and Torres 2010; Linos et al. 2009). Melanoma risk is increased by exposure to UVR from the sun and from tanning beds (Baldea et al. 2009; Wallingford et al. 2013).

Mutations in melanocytes arise from DNA damage caused either directly by UVB irradiation or indirectly following UVA irradiation through reactive oxygen (ROS) species via photosensitiser-mediated processes (Ikehata and Ono 2011). Solar UVR is composed of mainly UVA (320-400nm wavelengths) with a lesser component of UVB (280-320 nm wavelengths) and a UVA/UVB ratio of about 20 depending on latitude and time of day (Kollias et al. 2011). Melanocytes respond differently to UVA and UVB. UVA induces immediate pigment darkening through photo-oxidation of melanin that can be observed within minutes of exposure (BACHEM 1955; Hönigsmann et al. 1986). UVB on the other hand induces epidermal melanocyte proliferation (Nordlund et al. 1981; Rosdahl 1979; Sato and Kawada 1972) and activation of melanogenic enzymes resulting in a delayed tanning response occurring 2-3 days post exposure (BACHEM 1955). UVA and UVB also cause ROS mediated lipid peroxidation (Bose et al. 1990; Bose et al. 1989; Filipe et al. 2013), and widespread oxidative modification of proteins resulting in their proteasomal degradation (Grimm et al. 2012).

Raman spectroscopy enables a non-invasive label free analysis of the biochemical structures present within a sample (Brauchle et al. 2014; Moncada et al. 2016). Laser light is used to excite a sample and changes in scattering provide information about the molecular structures present (Gniadecka et al. 2004). Although previous studies have examined normal and melanoma biopsy tissue and compared melanocytes and melanoma cells (Brauchle et al. 2014; Silveira et al. 2012) none have examined melanocyte function, differentiation status or their UV response. Here, we use live single cell Raman spectroscopy and principal component analysis (PCA) to biochemically fingerprint melanoblasts, melanocytes, keratinocytes and melanoma cells. We identify the principal biomolecules that underlie these fingerprints and demonstrate that Raman spectroscopy is a highly sensitive method of probing the melanin biosynthesis pathway and its response to UVR.

## RESULTS

### Live single-cell Raman spectroscopy can discriminate between skin cell types and melanocyte differentiation status

We modelled keratinocytes, melanoblasts, melanocytes and melanoma cells using the COCA, melb-a, melan-a and B16F10 cell lines respectively (Figure 1a) (Diwakar et al. 2008; FIDLER 1973; Segrelles et al. 2011; Sviderskaya et al. 1995). Live single cell Raman spectroscopy was used to acquire individual spectra allowing us to establish their biochemical fingerprints. We confirmed viability of the cells after imaging using a CellTiter-Glo assay (Supplementary Figure 1a-c). The mean Raman spectra for each cell type (n = 9) appeared qualitatively distinct under normal culture conditions (Figure 1b). Principal component analysis (PCA) demonstrated that the main differences between the spectra were accounted for by cell type. We observed tight groupings for the melanoblasts, melanocytes and keratinocytes while melanoma cells were more heterogeneous in nature falling between the melanoblast and melanocyte groups on the PCA plot (Figure 1c). The PCA factor loadings (Figure 1d) were used to identify the principal Raman peaks responsible for the PCA groupings which we attributed to their likely biomarkers using the established literature (numbered 1-10 in Figure 1B and Supplementary Table 1). Nine of the ten peaks had been previously described and were consistent with the biochemical properties of the cell types examined including melanin biosynthesis (phenylalanine, tyrosine, melanin) photoprotection (β-carotene) or keratinocyte function (keratin/lipid) (Supplementary Table 1). We identified a peak at 1333 cm^-1^ specifically in melanocytes not previously attributed to a known biomarker but consistent with the melanin precursor DOPA (Supplementary Figure 1a and b).

**Figure 1.**
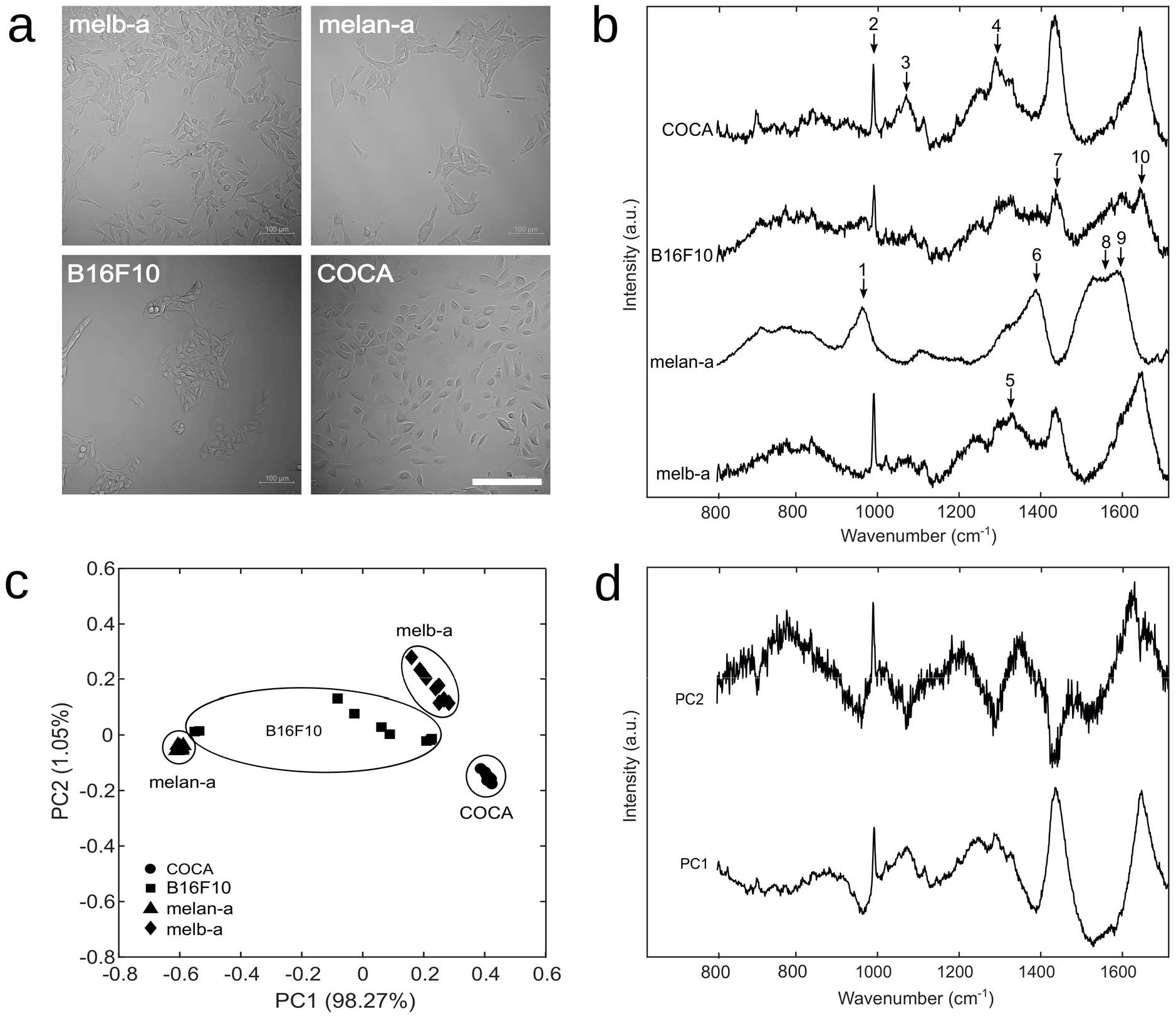
Live Raman spectroscopy can discriminate between cell types and differentiation status. Raman spectra (n = 9 for each cell type) were acquired from melanocytes (melan-a), melanoblasts (MelbA), melanoma cells (B16F10) and keratinocytes (COCA). Cells were grown on CaF_2_ disks in phenol red free medium prior to acquisition at 785nm. A principal component analysis (PCA) was then conducted. (a) Bright field images of the cell types used. (b) The mean Raman spectra (offset for clarity) for each cell type differed qualitatively. (c) A scatter plot of the principal component vectors PC1 (98.27%) and PC2 (1.05%) showing tight grouping of melanoblasts, melanocytes and keratinocytes with a more heterogeneous melanoma population. (d) Plot of the PC1 and PC2 vectors - the ten principal Raman peaks are labelled with arrows on (b). Scale bar in A = 200 μm. Numbers in (b): 1 = Lipid, 2 = Phenylalanine, 3 = Phospholipids/Tryptophan, 4 = Lipid, 5 = DOPA, 6 = β-carotene, 7 = Keratin, 8 = Melanin, 9 = Tyrosine, 10 = Amide I.

Undifferentiated melanoblasts and keratinocytes did not exhibit melanin peaks while the pigmented melanocytes and melanoma cells did (Figure 1b). Phenylalanine was apparent in melanoblasts, keratinocytes and melanoma cells but not in melanocytes presumably because melanin synthesis depletes the phenylalanine precursor in melanocytes. We observed DOPA peaks in melanoblasts and keratinocytes consistent with previous reports that they can convert tyrosine to DOPA (Gillbro et al. 2004; Schallreuter et al. 1995; Schallreuter et al. 1992).

### Immediate and temporal effects on melanin biosynthesis pathways after UV irradiation detected by Raman spectroscopy

Melanin biosynthesis proceeds through a stepwise conversion of phenylalanine through tyrosine and DOPA (Cichorek et al. 2013). As demonstrated above, melanocytes and melanoma cells are competent to perform all of these enzymatic steps while melanoblasts appear able to synthesise tyrosine but are unable to convert DOPA to melanin (Sviderskaya et al. 1995). Although the enzymatic pathways are well delineated the immediate and short-term biochemical response to UV exposure are still poorly understood. We therefore focused on the melanin biosynthesis pathway immediately after UV exposure by comparing log2 fold change (Log2-FC) in mean Raman peak height (n = 18 spectra in all cases) from control for the biomarkers identified above (Figure 2 and Table 1) in response to irradiation from a UVA, UVB and a mixed UVA/UVB light source over a one to 24-hour response period.

**Figure 2.**
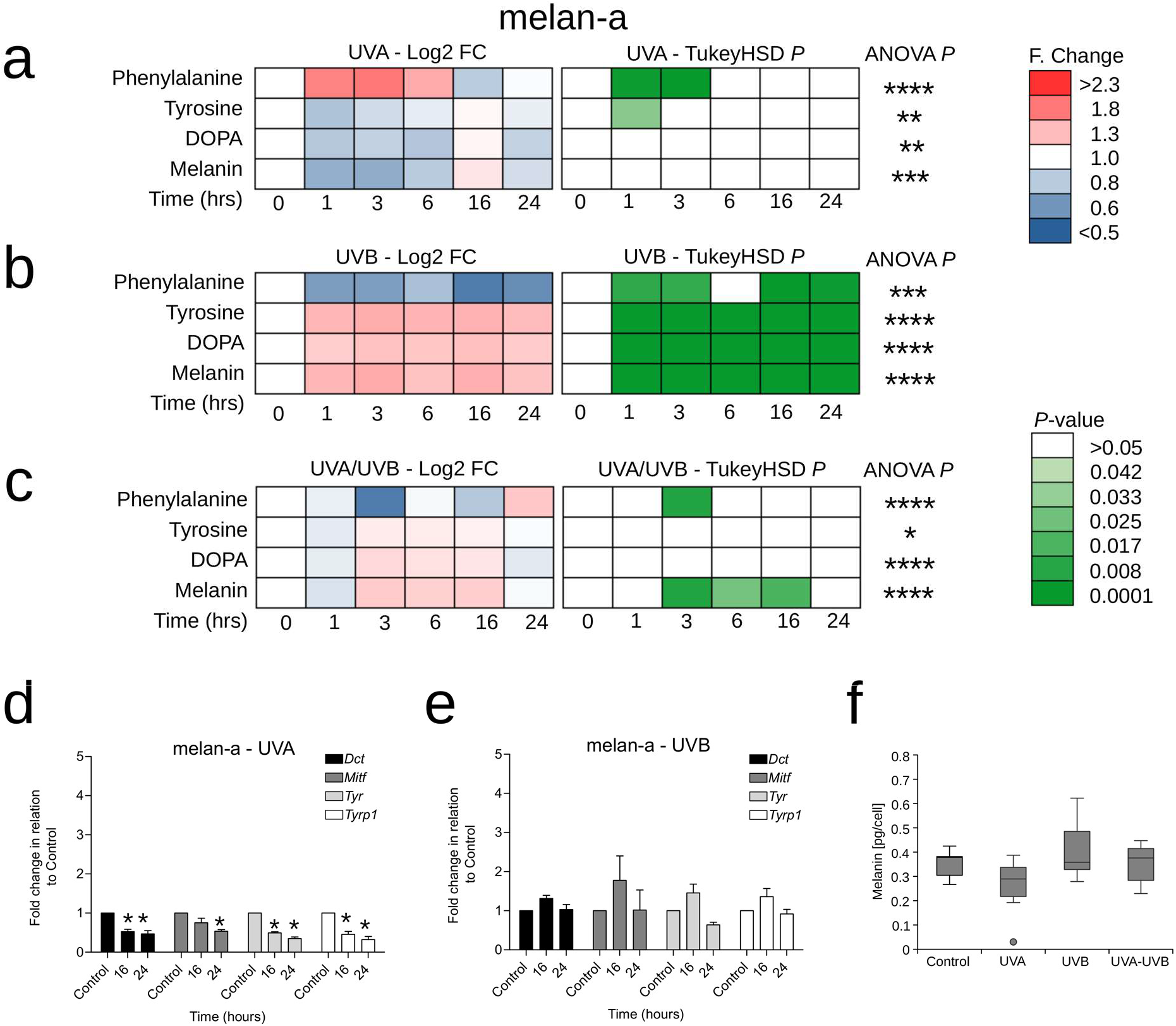
Distinct UVA and UVB responses in the melanin synthesis pathway in melan-a melanocytes. Cells were grown on CaF_2_ disks and irradiated with 100KJ/m^2^ UVA, 100J/m^2^ UVB or 1000J/m^2^ UVA + UVB. Raman spectra (at 785nm) were acquired at 1, 3, 6, 16 and 24 hours post irradiation. (a-c) Time course of log_2_ fold (F) change (Log2-FC) and pairwise Tukey’s honestly significant difference (TukeyHSD) comparing response (to UVA, UVB or UVA/UVB) to 0 hours control for phenylalanine, tyrosine, l-DOPA and melanin (n = 18 spectra for each square). Significance level for a one-way analysis of variance (ANOVA) is indicated at the end of each series (*<0.05, **<0.01, ***<0.001, ****<0.0001). (d-e) qRT-PCR time course analysis of fold change in *Tyr, Dct, Tyrp1* and *Mitf* expression (mean ± s.d, n=3) after. (f) Time course of intracellular melanin content (mean ± 95% CI, n = 3) after treatment.

### Opposing effects of UVA and UVB irradiation on the melanin biosynthesis pathway in melanocytes

Melanocytes were the most responsive to UVR. We observed a rapid (within one hour) and transient (up to six hours) statistically significant increase in the phenylalanine peak and a rapid (within one hour) and transient (up to six hours) statistically significant decrease in the tyrosine peak after UVA exposure (Figure 2a, Supplementary Figure 3a). In parallel we observed significant reductions in gene expression of *Mitf, Dct, Tyr* and *T*y*rp1* as measured by RT-qPCR at 16 and 24 hours post irradiation (Figure 2d). Together these results suggest a block in melanin synthesis and accumulation of the phenylalanine precursor in the 24 hours post UVA exposure. The changes in melanin levels detected by Raman after UVA irradiation were mirrored by similar trends in intracellular melanin measurement between 0 (control) and 1-3 hours (Figure 2f).

We observed opposing effects on the melanin biosynthesis pathway of UVB compared to UVA in melanocytes. This was characterised by a statistically significant rapid (within one hour) and sustained (up to 24 hours) decrease in the phenylalanine peak and a rapid (within one hour) and sustained (for up to 24 hours) statistically significant increase in the tyrosine, DOPA and melanin peaks (Figure 2b). Consistent with a rapid UVB response increasing melanin production independent of gene expression, we did not observe any accompanying changes in the gene expression levels of *Mitf, Dct, Tyr* and *T*y*rp1* as measured by RT-qPCR at 16 and 24 hours post irradiation (Figure 2e). The changes in melanin levels detected by Raman after UVB irradiation were mirrored by similar trends in intracellular melanin measurement between 0 (control) and 1-3 hours (Figure 2f).

The response of melanocytes to a mixed UVA/UVB light source appeared more nuanced with features of both the UVA and UVB responses (Figure 2c). However, the UVB response dominated with a rapid (one to three hours) and transient (up to three hours) statistically significant reduction in phenylalanine and a delayed (after one hour) and transient (less than 24 hours) statistically significant increase in the melanin peak mirrored by similar trends in intracellular melanin measurement between 0 (control) and 1-3 hours (Figure 2f).

### Similar effects of UVA and UVB irradiation on the melanin biosynthesis pathway in melanoma cells

While melanocytes appear to respond differently to UVA vs UVB, melanoma cells demonstrated similar responses to the three light sources. We observed a rapid (within one hour) and sustained (at least 24 hours) statistically significant increase in the phenylalanine peak post UVA irradiation (Figure3a) and a delayed (between six and 24-hours) and sustained (at least 24-hours) statistically significant increase in the phenylalanine peak post UVB and UVA/UVB irradiation (Figure 3b-c). This was accompanied by a statistically significant decrease in the melanin peak between one and 24-hours post UVB and UVA/UVB exposure (Figure 3b-c). The reduction in the melanin peak was the most pronounced after UVB treatment (Figure 3b) and we observed a corresponding reduction in the gene expression of *Tyrp1* at 16 and 24 hours suggesting a block in melanin synthesis (Figure 3e) similar to the one observed in melanocytes (melan-a) after UVA treatment above. The changes in melanin levels detected by Raman after UVA irradiation were mirrored by similar trends in intracellular melanin measurement between 0 (control) and 1-3 hours (Figure 2f).

**Figure 3.**
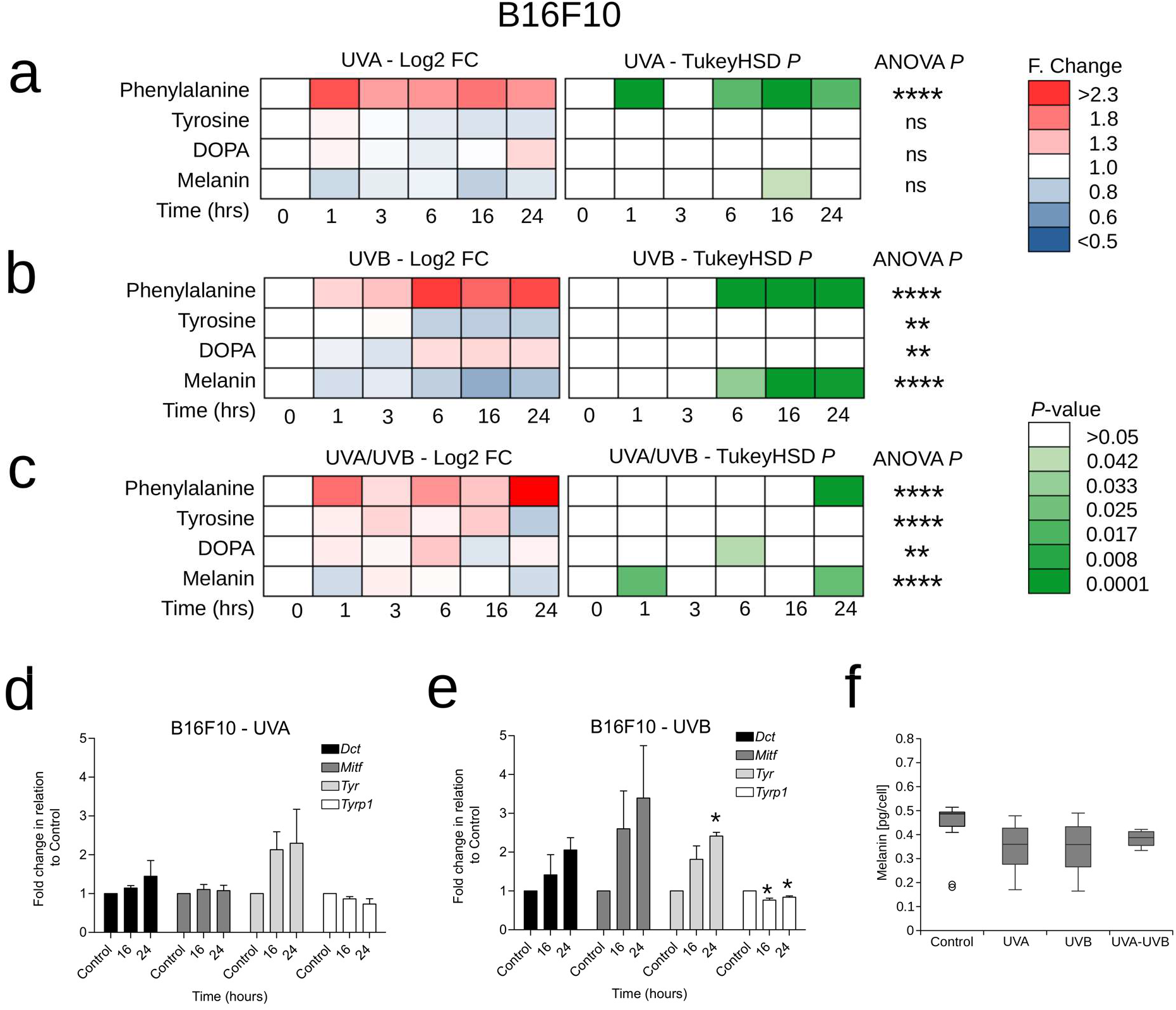
Similar UVA and UVB responses in the melanin synthesis pathway in B16F10 melanoma cells. Cells were grown on CaF2 disks and irradiated with 100KJ/m2 UVA, 100J/m2 UVB or 1000J/m2 UVA + UVB. Raman spectra (at 785nm) were acquired at 1, 3, 6, 16 and 24 hours post irradiation. (a-c) Time course of log_2_ fold (F) change (Log2-FC) and pairwise Tukey’s honestly significant difference (TukeyHSD) comparing response (to UVA, UVB or UVA/UVB) to 0 hours control for phenylalanine, tyrosine, l-DOPA and melanin (n = 18 spectra for each square). Significance level for a one-way analysis of variance (ANOVA) is indicated at the end of each series (*<0.05, **<0.01, ***<0.001, ****<0.0001). (d-e) qRT-PCR time course analysis of fold change in Tyr, Dct, Tyrp1 and Mitf expression (mean ± s.d, n=3) after. (f) Time course of intracellular melanin content (mean ± 95% CI, n = 3) after treatment.

### Minimal effects of UVA and UVB irradiation on the melanin biosynthesis pathway in melanoblasts and keratinocytes

We observed a subdued UV response in the melanin biosynthesis pathway in melanoblasts. Following UV exposure, we observed a delayed (between 16 and 24 hours) statistically significant reduction in the phenylalanine and the tyrosine peaks (Figure 4a) whilst following UVB exposure we saw an immediate (between 1 and 3 hours) and transient (up to 6 hours) statistically significant reduction in the phenylalanine and tyrosine peaks (Figure 4b). The response to the mixed UVA/UVB light source had features of the UVA and UVB responses with immediate (within one hour) and delayed (by 16-24 hours) statistically significant reductions in the phenylalanine and tyrosine peaks (Figure 4c). Taken with the results from melanocytes and melanoma cells these results suggest that whilst accumulation of phenylalanine may be due to a block in the melanin biosynthesis pathway, reductions in phenylalanine, detected by Raman spectroscopy, can occur both through rapid production of melanin depleting the phenylalanine precursor but may also occur independently perhaps through photodegradation as previously reviewed (Kerwin and Remmele 2007).

**Figure 4.**
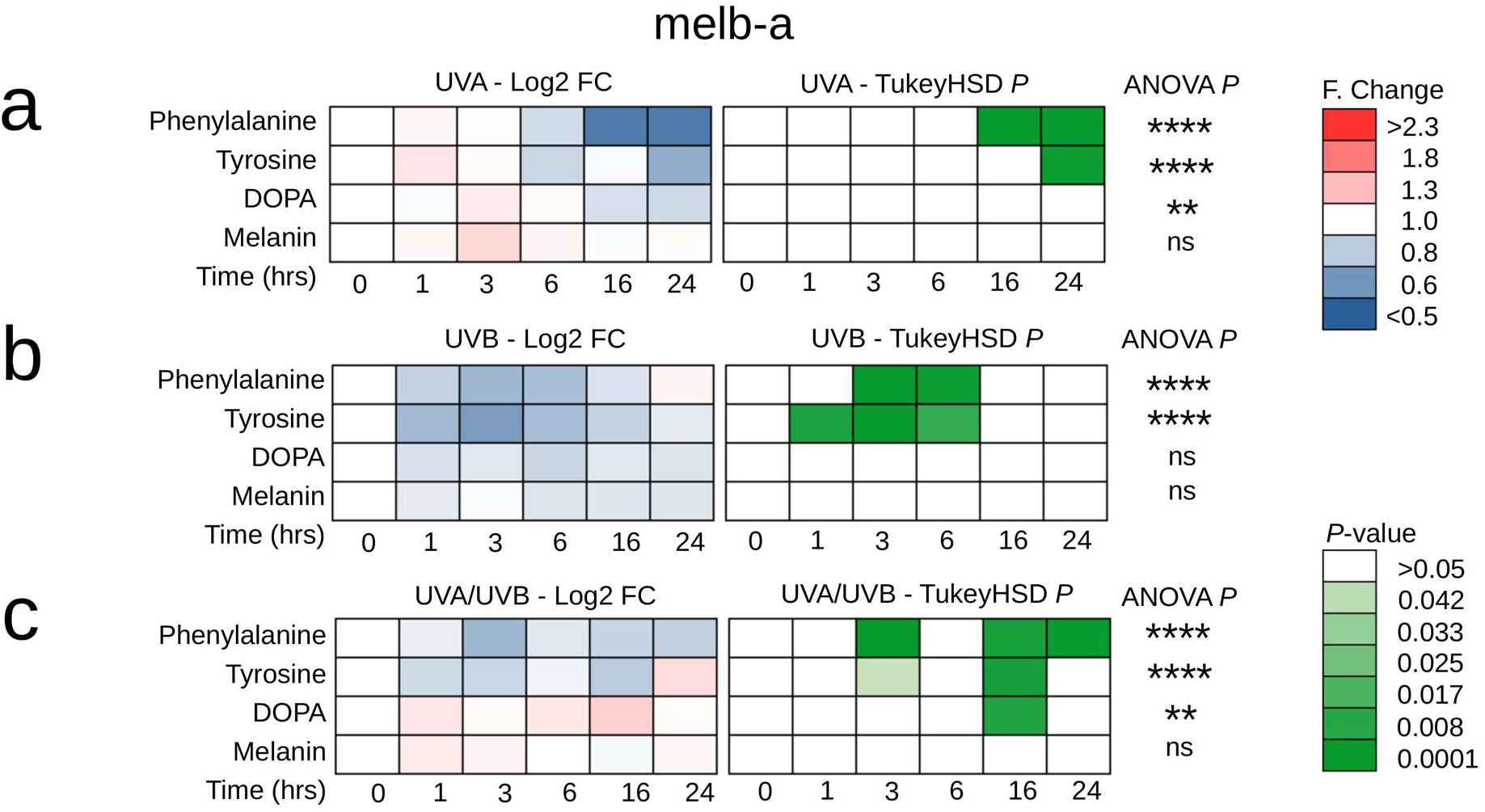
UVA and UVB responses in the melanin synthesis pathway in melanoblasts (melb-a). Cells were grown on CaF_2_ disks and irradiated with 100KJ/m2 UVA, 100J/m2 UVB or 1000J/m2 UVA + UVB. Raman spectra (at 785nm) were acquired at 1, 3, 6, 16 and 24 hours post irradiation. (a-c) Time course of log_2_ fold (F) change (Log2-FC) and pairwise Tukey’s honestly significant difference (TukeyHSD) comparing response (to UVA, UVB or UVA/UVB) to 0 hours control for phenylalanine, tyrosine, l-DOPA and melanin (n = 18 spectra for each square). Significance level for a one-way analysis of variance (ANOVA) is indicated at the end of each series (*<0.05, **<0.01, ***<0.001, ****<0.0001).

Consistent with activity of the melanin biosynthesis pathway downstream of tyrosinase (TYR) being the key determinant in the Raman spectra observed in cells of the melanocyte lineage we observed a subdued UV response in keratinocytes very similar to the one observed in undifferentiated melanoblasts with a delayed (24 hours) statistically significant reduction in phenylalanine post UVA exposure (Figure 5a) and an immediate (from one hour) and sustained (up to 24-hours) reduction in phenylalanine and tyrosine post UVB exposure (Figure 5b). Exposure to the UVA-UVB mixed light source resulted in an immediate (one hour) and transient (up to three hours) reduction in tyrosine (Figure 5c). We observed inconsistent changes in the DOPA and melanin peaks in keratinocytes likely attributable to noise caused by the very low basal level of DOPA in the control samples (Figure 5a).

**Figure 5.**
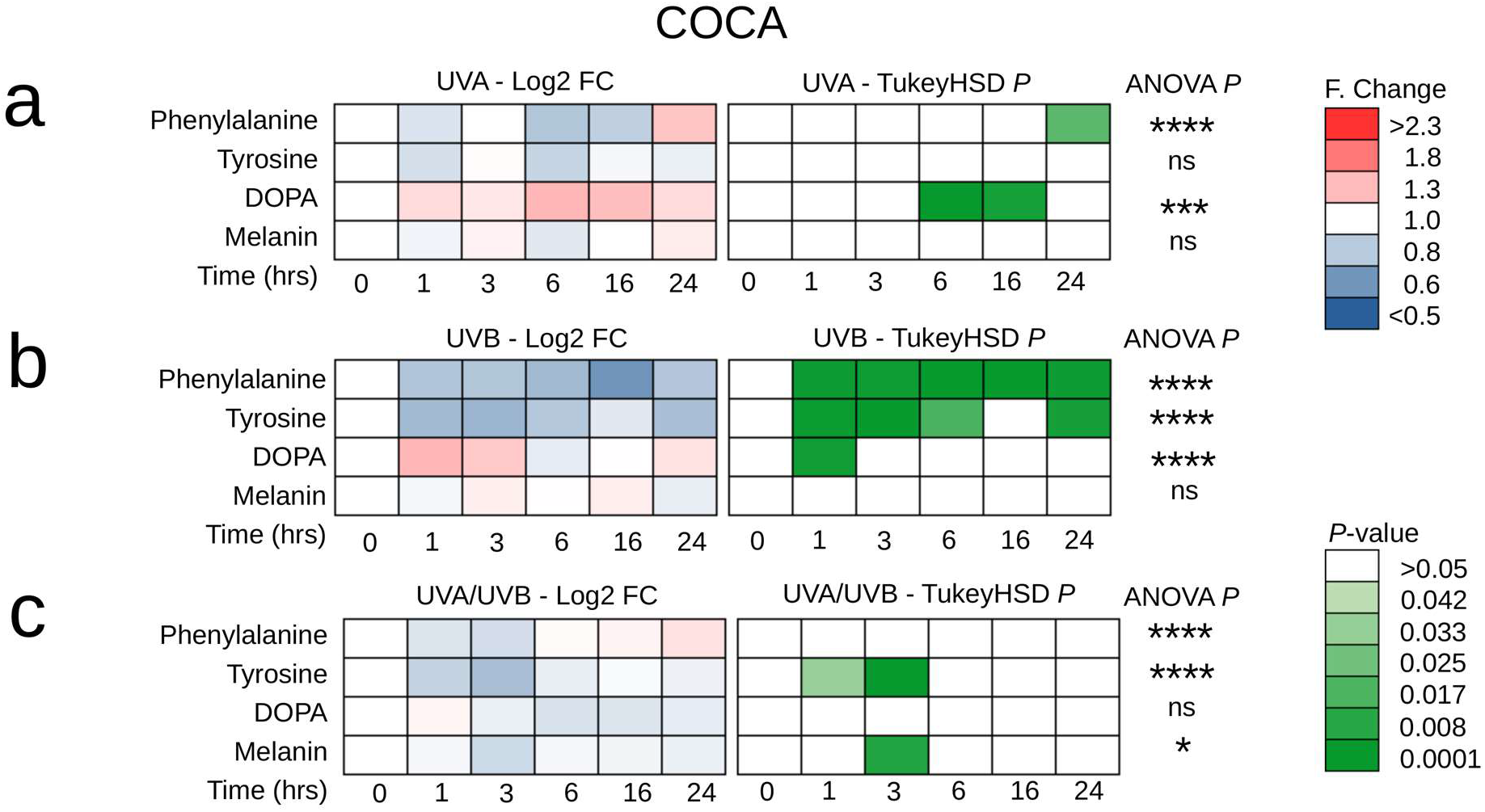
UVA and UVB responses in the melanin synthesis pathway in keratinocytes (COCA). Cells were grown on CaF2 disks and irradiated with 100KJ/m2 UVA, 100J/m2 UVB or 1000J/m2 UVA + UVB. Raman spectra (at 785nm) were acquired at 1, 3, 6, 16 and 24 hours post irradiation(a-c) Time course of log_2_ fold (F) change (Log2-FC) and pairwise Tukey’s honestly significant difference (TukeyHSD) comparing response (to UVA, UVB or UVA/UVB) to 0 hours control for phenylalanine, tyrosine, l-DOPA and melanin (n = 18 spectra for each square). Significance level for a one-way analysis of variance (ANOVA) is indicated at the end of each series (*<0.05, **<0.01, ***<0.001, ****<0.0001).

### Differentiated melanocytes but not melanoblasts, melanoma cells or keratinocytes demonstrate a photoprotective responses to UVR

Melanocytes have been shown to preferentially absorb β-carotene in culture (Andersson et al. 2001). Consistent with this, we observed a distinct β-carotene Raman peak only in melanocytes (Figure 1b, Supplementary Table 1). In response to UVR we observed a statistically significant increase in this β-carotene peak (between one and 24 hours post UV) in response to UVA, UVB and mixed UVA/UVB light sources (Figure 6b). We therefore hypothesised that β-carotene may be performing a cell-type specific photoprotective function protecting proteins and lipids from oxidative damage. We investigated changes in the Raman peaks for lipids (1300cm^-1^ peak) and amide I (protein peaks at 1650cm^-1^) in our Raman spectra as biomarkers of oxidative damage (Figure 6a-d).

**Figure 6.**
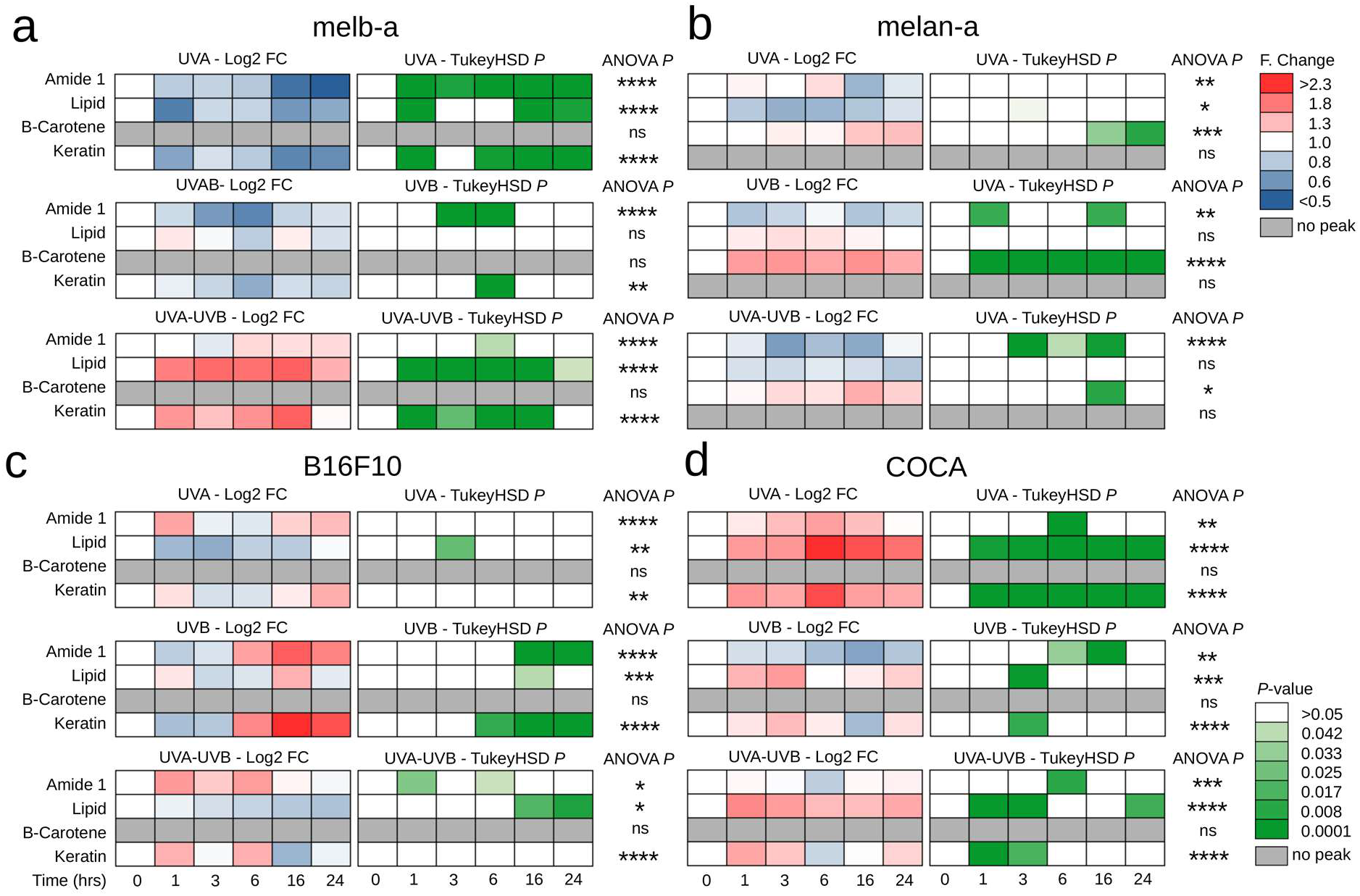
Heterogeneous responses of melanoblasts, melanocytes, melanoma cells and keratinocytes to UVR. Cells were grown on CaF2 disks and irradiated with 100KJ/m2 UVA, 100J/m2 UVB or 1000J/m2 UVA + UVB. Raman spectra (at 785nm) were acquired at 1, 3, 6, 16 and 24 hours post irradiation. (a-d) Time course of log2 fold (F) change (Log2-FC) and pairwise Tukey’s honestly significant difference (TukeyHSD) test comparing response to 0 hours control for amide I, lipid, keratin, and β-carotene (n = 18 spectra for each square). Significance level for a one-way analysis of variance (ANOVA) is indicated at the end of each series (*<0.05, **<0.01, ***<0.001, ****<0.0001).

We observed a rapid (one to three hours) and statistically significant reduction in the amide I peak in melanoblasts in response to UVA and UVB (Figure 6a) which appeared absent or dampened in melanocytes (Figure 6b). Furthermore, we observed a rapid (within one hour) statistically significant reduction in the lipid peak in response to UVA and a rapid (within one hour) statistically significant increase in the lipid peak in response to a mixed UVA/UVB light source in melanoblasts (Figure 6a). No changes in the lipid peak were observed in melanocytes (Figure 6b). These results suggest differentiated melanocytes are better protected against the effects of UVR than melanoblasts. Consistent with a more heterogeneous population of cells in melanoma, no clear trends in the lipid or amide I peaks were observed in melanoma cells (Figure 6c). Lipid levels have previously been shown to increase in human keratinocytes post UVA and UVB dual irradiation (Olivier et al. 2017). Consistent with this, here we observed rapid (from 1-3 hours) and sustained (at least 24 hours) statistically significant increases in the lipid peak in response to UVA, UVB and our mixed UVA/UVB light source in keratinocytes (Figure 6d).

### Melanoblasts, melanoma cells and keratinocytes but not melanocytes demonstrate changes in keratin levels on UV exposure

Raman peaks consistent with keratin were present in keratinocytes and melanoblasts. We observed a rapid (within 1 hour) and sustained (up to 24 hours) statistically significant increase in the Raman peak for keratin in keratinocytes after UVA treatment with similar rapid (one to three hours) and transient (less than six hours) statistically significant increases after UVB and combined UVA/UVB treatments in these cells (Figure 6d). Melanoblasts displayed a significant reduction in keratin after individual UVA and UVB treatments but an immediate (from one hour) and sustained (up to 24 hours) statistically significant increase in keratin after exposure to the dual UVA/UVB light source (Figure 6a). Melanoma cells but not melanocytes also exhibited a keratin peak and demonstrated a delayed (6-24 hours) statistically significant increase after UVB treatment (Figure 6c).

## DISCUSSION

We demonstrate the utility of live Raman spectroscopy performed on cells grown in a controlled environmental chamber, as a unique tool to probe the melanin biosynthesis pathway and its immediate response to UVR and reveal rapid and opposing responses to UVA and UVB irradiation by melanocytes.

### Raman fingerprinting demonstrates that melanoma cells are phenotypically heterogeneous

Previous reports have discriminated between malignant melanoma cells and their wildtype counterparts using Raman spectroscopy (Silveira Jr. et al., 2012; Brauchle et al., 2014). Our Raman spectra appear consistent with those previously reported for melanocytes and melanomas cells (Feng et al. 2017; Gniadecka et al. 2004; Silveira et al. 2012), while our spectra for melanoblasts and keratinocytes are consistent with their inability to synthesise melanin (Sviderskaya, Wakeling and Bennett, 1995). Our PCA analysis demonstrates that melanoblasts, melanocytes and keratinocytes represent biochemically homogeneous populations while melanoma cells are a biochemically heterogeneous positioned between undifferentiated melanoblasts and differentiated melanocytes. This is consistent with their known phenotypic heterogeneity and the reactivation of their embryonic transcriptome (Marie et al. 2020; Vandamme and Berx 2014). Further study is warranted to address whether this method can be used to predict the position of a melanoma cell on the differentiation trajectory of melanoma subtypes (Tsoi et al. 2018) which may allow exploitation of these properties to increase immune therapy efficacy.

### Rapid and opposing effects of UVA and UVB irradiation on the melanin biosynthesis pathway in melanocytes

We show here the utility of Raman spectroscopy to detect and sensitively measure changes in the melanin biosynthesis pathway in response to UVR over a smaller time-frame than is possible with conventional methods. UVA exposure has long been understood to result in immediate pigment darkening within a few hours of exposure by acting on pre-existing melanin and by *de novo* melanin synthesis (Miyamura et al. 2011; Wicks et al. 2011). The reduction in melanin post UVA and accumulation of phenylalanine observed here may be consistent with a rapid tanning response resulting in the export of melanosomes from the cytosol and a subsequent block in synthesis because of their loss leading to accumulation of phenylalanine, consistent with reports UVA induces a redistribution of melanosomes in the skin (Lavker and Kaidbey 1982). However we do not see the rapid accumulation of melanin overserved in human epidermal melanocytes (Wicks et al. 2011). UVB has long been demonstrated to cause a delayed tanning response mediated through DNA damage induced *de novo* tyrosinase production (Cui et al. 2007; Eller et al. 1996). We were surprised to see such a rapid and profound UVB mediated response in the present study characterised by rapid and sustained increases in tyrosine, DOPA and melanin and reductions in phenylalanine suggesting that a previously overlooked mechanism independent of DNA damage acts immediately after UVB exposure in mouse melanocytes.

### Melanocytes uniquely harbour β-carotene with a possible role in photoprotection

The photoprotective effects of β-carotene have been attributed to its free radical/reactive oxygen species (ROS) scavenging abilities as well as direct absorption of UV at 400 nm (Alaluf et al. 2002; Moshell and Bjornson 1977). We observe melanocyte specific accumulation of β-carotene consistent with previous reports (Andersson, Vahlquist and Rosdahl, 2001) and report here that β-carotene levels increase significantly in response to UVA, UVB and a mixed UVA/UVB light source. The presence of β-carotene also appeared to correlate with an apparent stabilisation in the peaks of amide I and lipid in melanocytes, whereas these biomarkers appeared to change dramatically in response to UVR in the other cell types studied. Taken together these finding suggest that melanocytes use β-carotene as part of a UVR protective mechanism shielding against UV-induced lipid peroxidation, DNA damage and protein oxidation as previously reported (Alaluf et al., 2002; Stephen, Coetzee and Perrett, 2011).

## CONCLUSION

We have defined a reference biochemical signature for each of the cell types studied that can be attributed to their biological function. In doing so we have demonstrated the utility of Raman spectroscopy as a tool with which to probe the melanin biosynthesis pathway with greater temporal resolution and sensitivity than conventional methods. We have used this approach to uncover hitherto undescribed immediate responses to UVA and UVB irradiation in melanocytes including a possible photoprotective role for β-carotene. The ability to probe biochemical status with high temporal resolution will be highly beneficial to continued research in the study of melanoma and the further development of sensitive screening and early detection methods.

## MATERIALS AND METHODS

### Cell culture

Mouse melan-a cells were cultured in RPMI 1640 (21875-034, Invitrogen) supplemented with 10% (v/v) FCS and 200nM TPA. melb-a cells were cultured in RPMI 1640 (21875-034, Invitrogen) supplemented with 10% (v/v) FCS, 40pM FGF2 and 20ng/ml mSCF. B16F10 cells were cultured in RPMI 1640 (21875-034, Invitrogen) supplemented with 5% (v/v) FCS. COCA cells were cultured in CnT-07 (CELLnTEC). Cells were incubated at 37°C in humidified air containing 5% (v/v) CO_2_.

### Raman spectroscopy

Cells were seeded onto CaF_2_ disks in 24-well plates using phenol-free media. Cells were irradiated with 100KJ/m2 UVA, 1000J/m^2^ UVA and UVB or 100J/m2 UVB and spectra acquired at 1, 3, 6, 16 and 24 hours post radiation as well as from an untreated control. Raman spectra were acquired at 60x using a 785nm laser at a spectral range of 600-1700cm^-1^ using an InVia Raman microspectrometer (Renishaw plc, Gloucestershire, UK) equipped with an environmental chamber (Okolab, Ottaviano, NA, Italy) maintaining a temperature of 37°C in humidified air containing 5% (v/v) CO_2_. Collection was at 2 seconds with 9 accumulations (18 seconds total) at 100% laser power. A total of 18 spectra were collected across 3 independent runs with spectra from 6 individual cells collected each run. Cells were selected randomly by moving diagonally along the sample and collecting spectra every 10^th^ cell to ensure spread across the disc and that no overlap occurred. Data collected was baseline corrected using a polynomial method order 3, smoothed using P=0.001, λ=105 then vector normalised. Principal component analysis was performed in MatlabR2018A (The Mathworks, NA, USA) using an in-house script (Martin et al. 2010; Trevisan et al. 2013).

### Cell Viability

Viability was measured using Cell Titer Glo (promega). CaF_2_ disks were transferred to a 24 well plate, along with control disks that had not been used for spectral acquisition. Diluted Cell Titer Glo was added to cells for 10 minutes before scraping and transfer to a white 96 well plate. Bioluminescence was measured as relative light units using a Perkin Elmer Wallac 1420 Victor2 microplate reader.

### Intracellular melanin measurements

Cells were UV-irradiated as described above. Subsequently they were washed in PBS, trypsinised and counted, then pelleted at 1000 RPM for 3 minutes. 100μL of melanin lysis buffer (90% 1M NaOH and 10% DMSO) was added to each pellet before incubation at 80°C for 90 minutes. Lysed pellets were transferred to a 96 well plate and their absorbance read at 490nm on a Perkin Elmer Wallac 1420 Victor2 microplate reader. A standard curve was obtained by diluting synthetic melanin (Sigma, Poole, UK) at 0 to 1μg/mL to enable the final melanin concentration to be determined.

### Statistics

All statistical tests were performed in SPSS or using the ‘R’ statistics package (http://www.R-project.org). The Raman data in Figures 2-6, was normalised individually for each peak across all cell types and conditions. A one-way analysis of variance (ANOVA) was performed across the six time points for each combination of cell type and UVA treatment followed by pairwise comparisons by Tukey’s Honestly Significant Difference test (TukeyHSD). Therefore both a significant ANOVA *P* value and a significant TukeyHSD *P* value is required for a significant pairwise difference to be considered. For the qPCR data in Figures 2 and 3, data was analysed using SPSS. A one-way analysis of variance (ANOVA) was performed across the 3 time points for each cell type and UVR treatment this was followed by pairwise Dunnett’s post hoc tests. Significant differences between control and time were evaluated with *p values ≤0.05 as indicated on the graphs.

### RT-qPCR Analysis

Total RNA was extracted using the RNAeasy kit (Qiagen, Crawley, UK) following the manufacturer’s instructions. Reverse transcription of 1 μg RNA was performed using oligo dT18, M-MLV reverse transcriptase and RNase OUT (Invitrogen, Thermo Fisher Scientific) to generate cDNA. For RT-qPCR 10ng cDNA was used along with 12.5 μl of 2X Power SYBR® Green master mix (Applied Biosystems, Warrington, UK) and 400nM primers (Sigma, Poole, UK). Reactions were performed on a BioRad CFX RT-qPCR machine using the following parameters: 50°C for 2 min and 95°C for 10 min followed by 40 cycles of 95°C for 15 seconds and 60°C for 1 min. Cycle threshold (CT) values were calculated for each mRNA sample and compared to their respective actin control to determine gene expression changes using the comparative CT (2−ΔΔCT) method (Livak and Schmittgen 2001).

## DATA AVAILABILITY STATEMENT

The data that support the findings of this study will be available after publication at https://www.research.lancs.ac.uk/portal/en/people/richard-mort(ed9f7471-68c9-4816-97a9-58f8f44c1f1d).html and from the corresponding author on reasonable request.

## ACKNOWLEDGMENTS

This work was supported by a project grant from North West Cancer Research (Grant CR1132).

## AUTHOR CONTRIBUTIONS STATEMENT

Conceptualization: RLM, SLA, JGK, LA; Data Curation: RLM, EW, JGK; Formal Analysis: ELW, RLM; Funding Acquisition: RLM, SLA; Investigation: ELW; Methodology: RLM, JGK; Project Administration: ELW; Resources: RLM, SLA, JGK, LA; Software: JGK, RLM, ELW; Supervision: RLM; Validation: ELW; Visualization: ELW, RLM; Writing - Original Draft Preparation: ELW, RLM; Writing - Review and Editing: RLM, ELW, SLA, LA, JGK.

## GRAPHICAL ABSTRACT

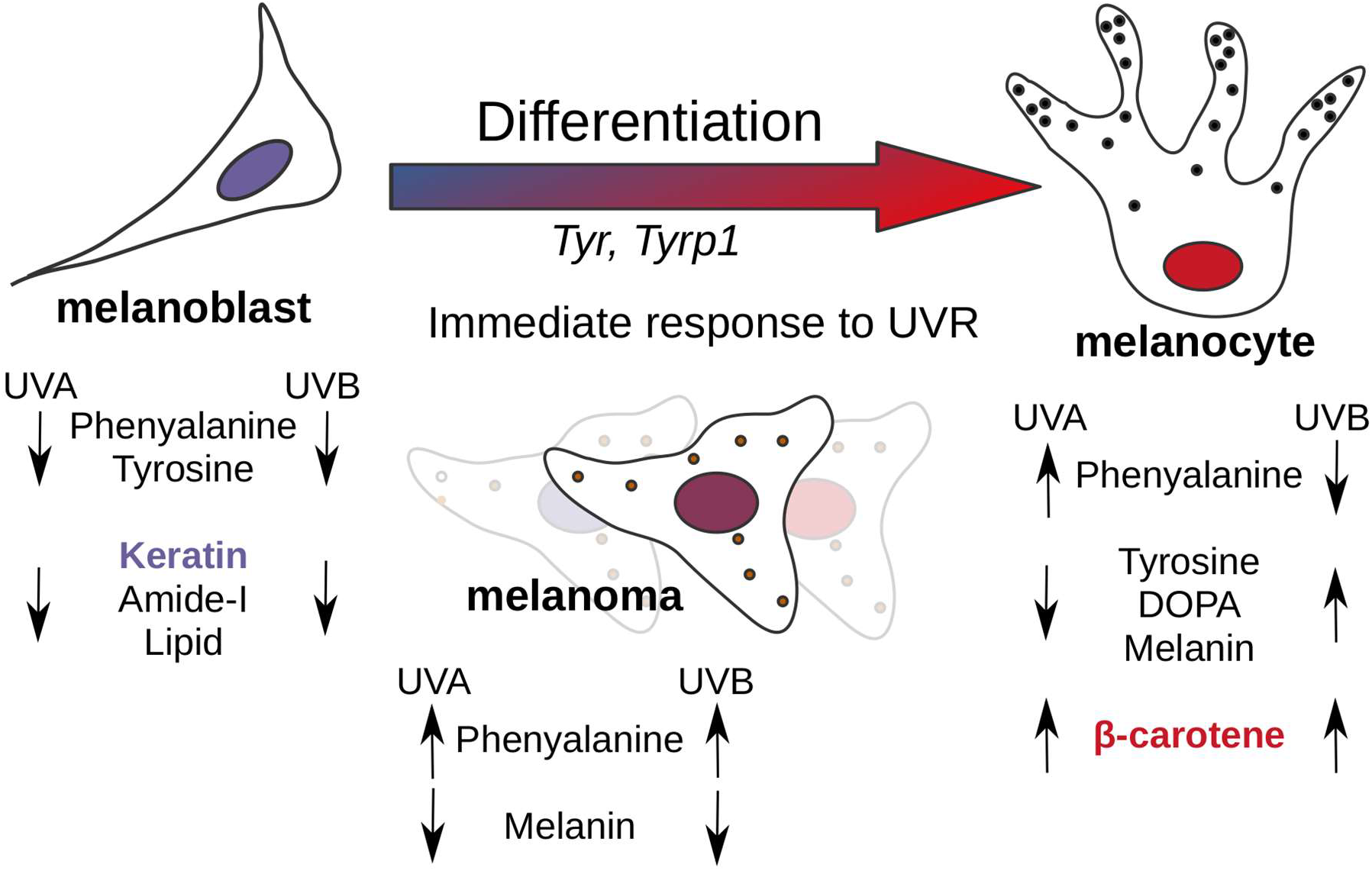

## SUPPLEMENTARY MATERIAL

## SUPPLEMENTARY TABLES

**Supplementary Table 1:**
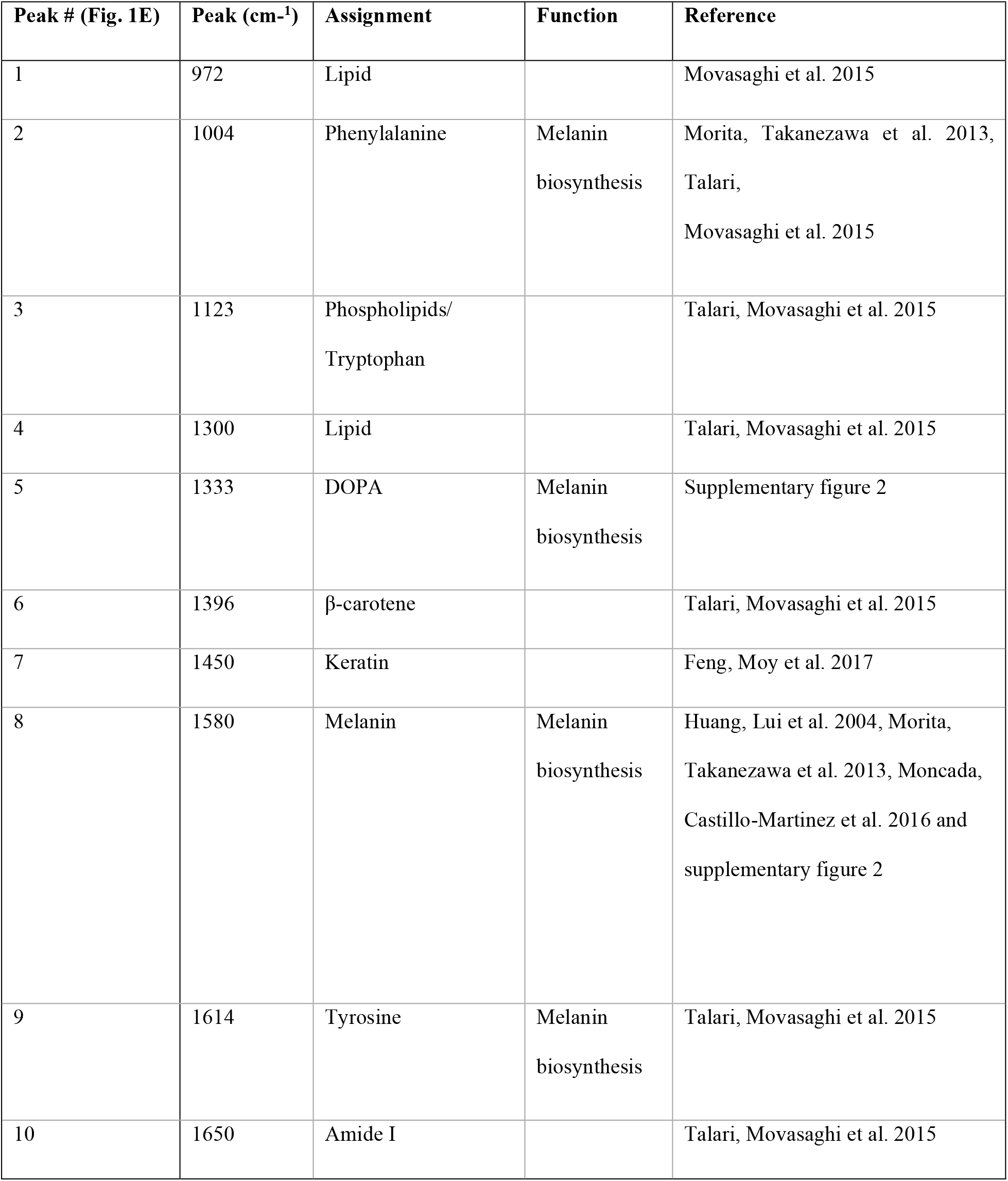
Summary of Raman spectral peaks.

**Supplementary Table 2:**
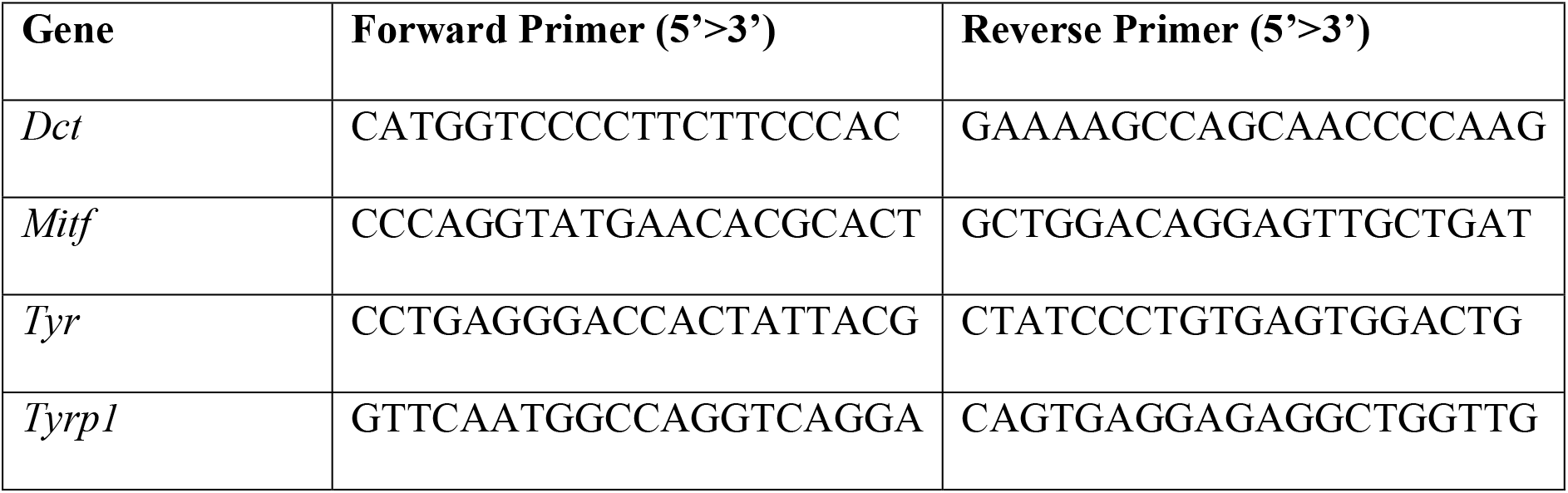
Oligonucleotides used for RT-qPCR analysis.

## SUPPLEMENTARY FIGURES

**Supplementary Figure 1.**
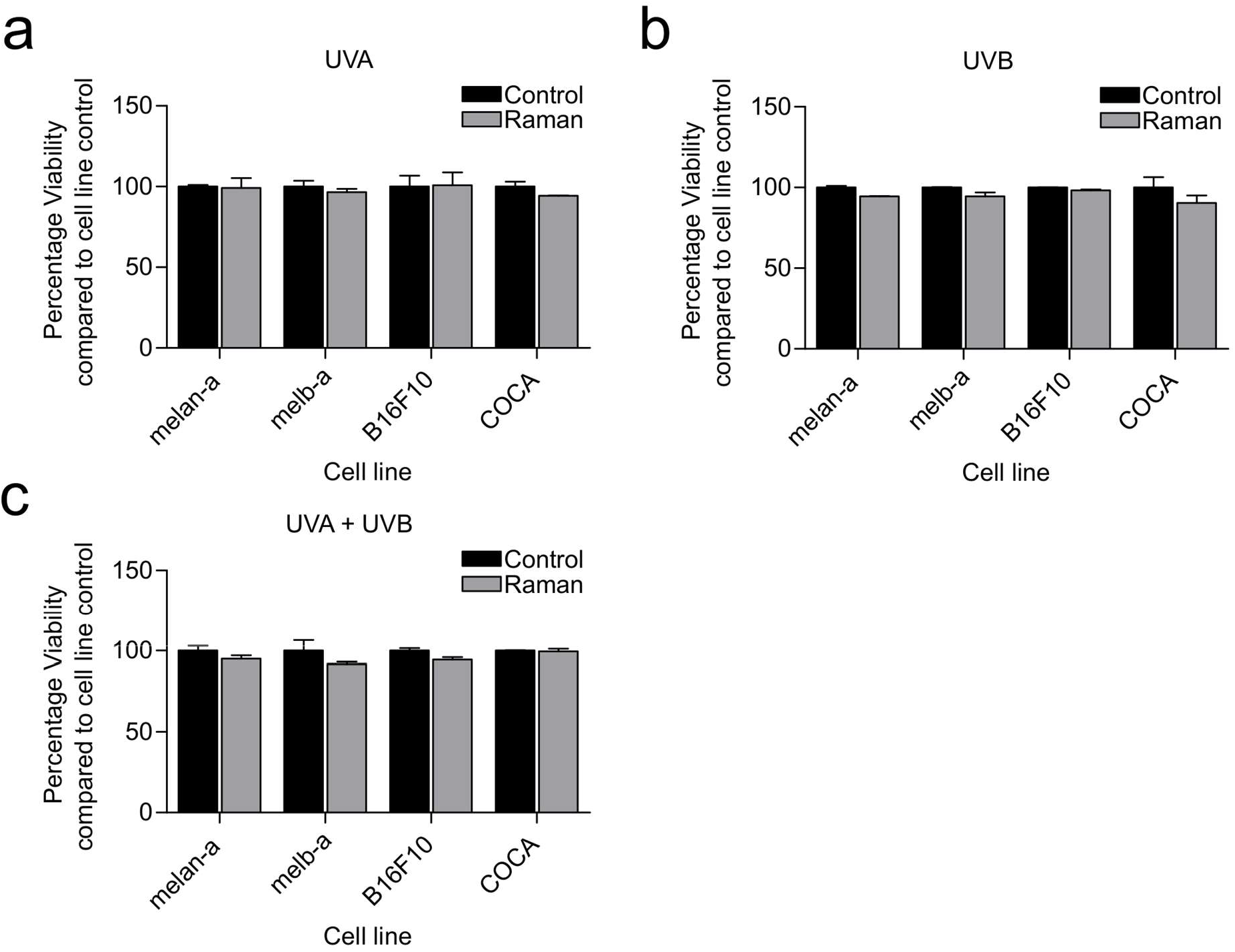
Cell viability post Raman spectral acquisition. melan-a, melbA, B16F10 and COCA cells were grown on CaF_2_ disks in duplicate in phenol red free medium for 24 hours before irradiation with 100KJ/m^2^ UVA, 1000J/m^2^ UVA and UVB or 100J/m^2^ UVB before spectra acquired from one disk at 1, 3, 6, 16 and 24 hours post radiation as well as from an untreated control. The other disk was maintained under normal culture conditions as a control. Cell viability was measured using Cell Titer Glo. (A) UVA, (B) UVB and (C) UVA and UVB. Data represented as mean ± 95% CI, N=18.

**Supplementary Figure 2.**
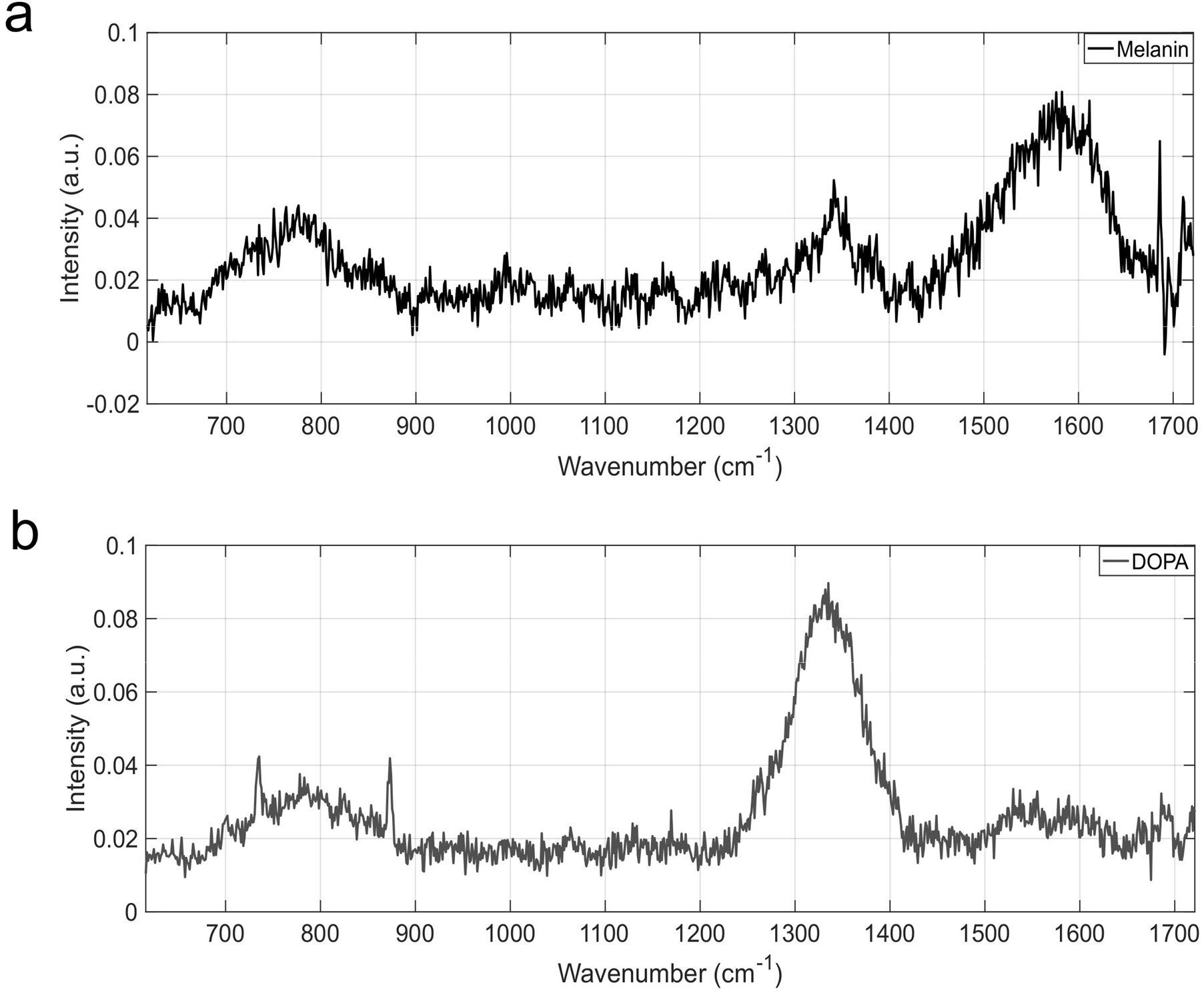
Raman spectra of synthetic melanin and DOPA. The spectra of melanin and DOPA was acquired using a 785nm laser at a spectral range of 600-1700cm^-1^. Spectra was collected at 2 seconds with 9 accumulations (18 seconds total) at 100% laser power. Data collected was baseline corrected and smoothed using P=0.001, λ=105 before vector normalising.

**Supplementary Figure 3.**
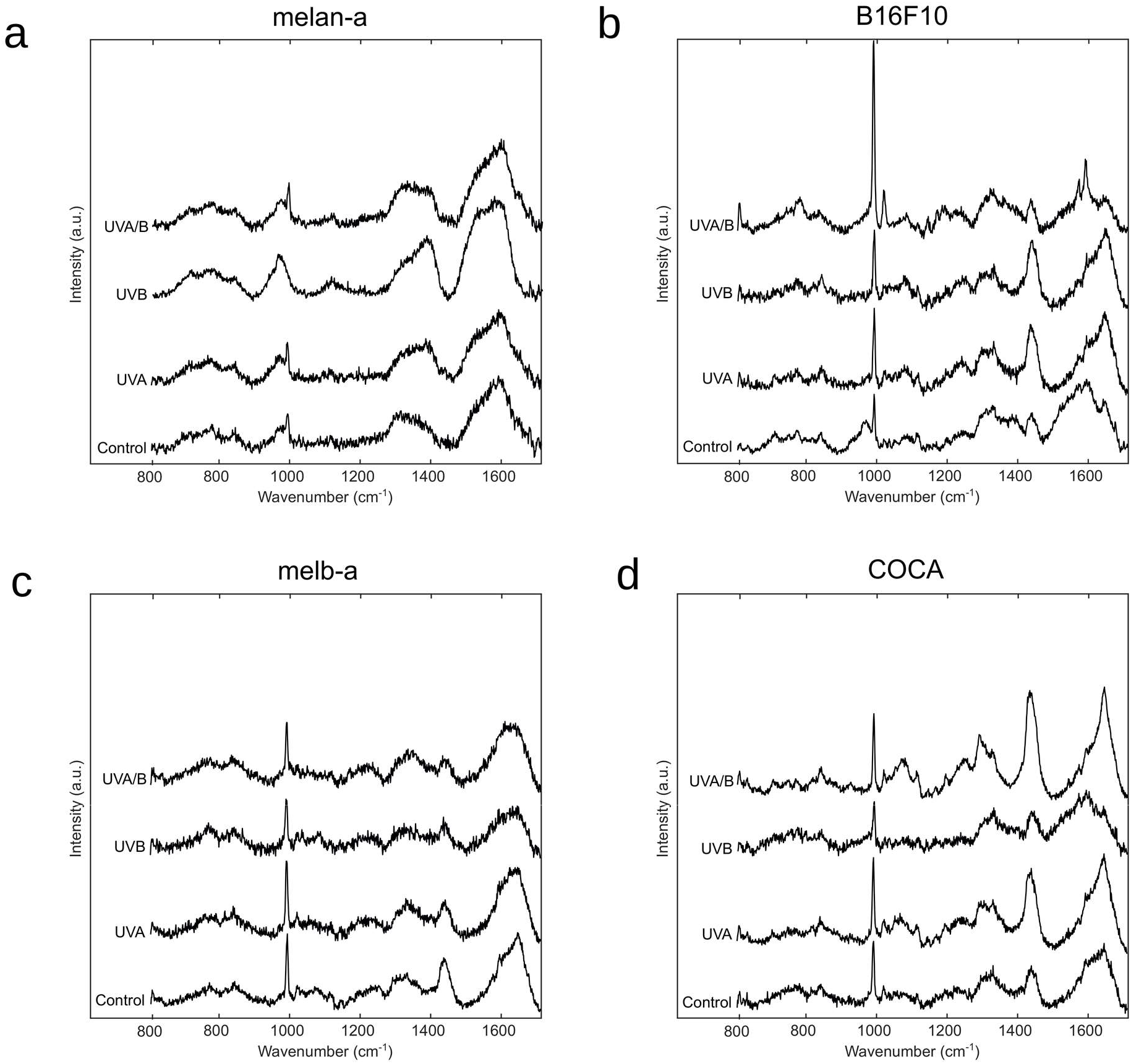
Raman spectra of UVR treated cells. melan-a, melbA, B16F10 and COCA cells were grown on CaF_2_ disks and irradiated with 100KJ/m^2^ UVA, 1000J/m^2^ UVA and UVB or 100J/m^2^ UVB before Raman spectra acquired using a 785nm laser at a spectral range of 600-1700cm^-1^, 100% laser power 2 seconds with 9 accumulations (18 seconds total) at 24 hours post irradiation and compare to an untreated control. Data collected was baseline corrected and smoothed using P=0.001, λ=105 before vector normalising. Raman spectra for each cell line plotted as class mean (n=18) (A) melan-a, (B) B16F10, (C) melbA, (D) COCA.

