## Supplementary material for "Fingerprinting of skin cells by live cell Raman spectroscopy reveals melanoma cell heterogeneity and cell-type specific responses to UVR": Supplemntal Material

### SUPPLEMENTARY TABLES:

#### Supplementary Table 1: Summary of Raman spectral peaks.

| Peak # (Fig. 1E) | Peak ($cm^{-1}$) | Assignment | Function | Reference |
|---|---|---|---|---|
| 1 | 972 | Lipid | | Movasaghi et al. 2015 |
| 2 | 1004 | Phenylalanine | Melanin biosynthesis | Morita, Takanezawa et al. 2013, Talari, Movasaghi et al. 2015 |
| 3 | 1123 | Phospholipids/ Tryptophan | | Talari, Movasaghi et al. 2015 |
| 4 | 1300 | Lipid | | Talari, Movasaghi et al. 2015 |
| 5 | 1333 | DOPA | Melanin biosynthesis | Supplementary figure 2 |
| 6 | 1396 | β-carotene | | Talari, Movasaghi et al. 2015 |
| 7 | 1450 | Keratin | | Feng, Moy et al. 2017 |
| 8 | 1580 | Melanin | Melanin biosynthesis | Huang, Lui et al. 2004, Morita, Takanezawa et al. 2013, Moncada, Castillo-Martinez et al. 2016 and supplementary figure 2 |
| 9 | 1614 | Tyrosine | Melanin biosynthesis | Talari, Movasaghi et al. 2015 |
| 10 | 1650 | Amide I | | Talari, Movasaghi et al. 2015 |

**Supplementary Table 2: Oligonucleotides used for RT-qPCR analysis.**

| **Gene** | **Forward Primer (5’>3’)** | **Reverse Primer (5’>3’)** |
|---|---|---|
| *Dct* | CATGGTCCCCTTCTTCCCAC | GAAAAGCCAGCAACCCCAAG |
| *Mitf* | CCCAGGTATGAACACGCACT | GCTGGACAGGAGTTGCTGAT |
| *Tyr* | CCTGAGGGACCACTATTACG | CTATCCCTGTGAGTGGACTG |
| *Tyrp1* | GTTCAATGGCCAGGTCAGGA | CAGTGAGGAGAGGCTGGTTG |

**SUPPLEMENTARY FIGURES:**

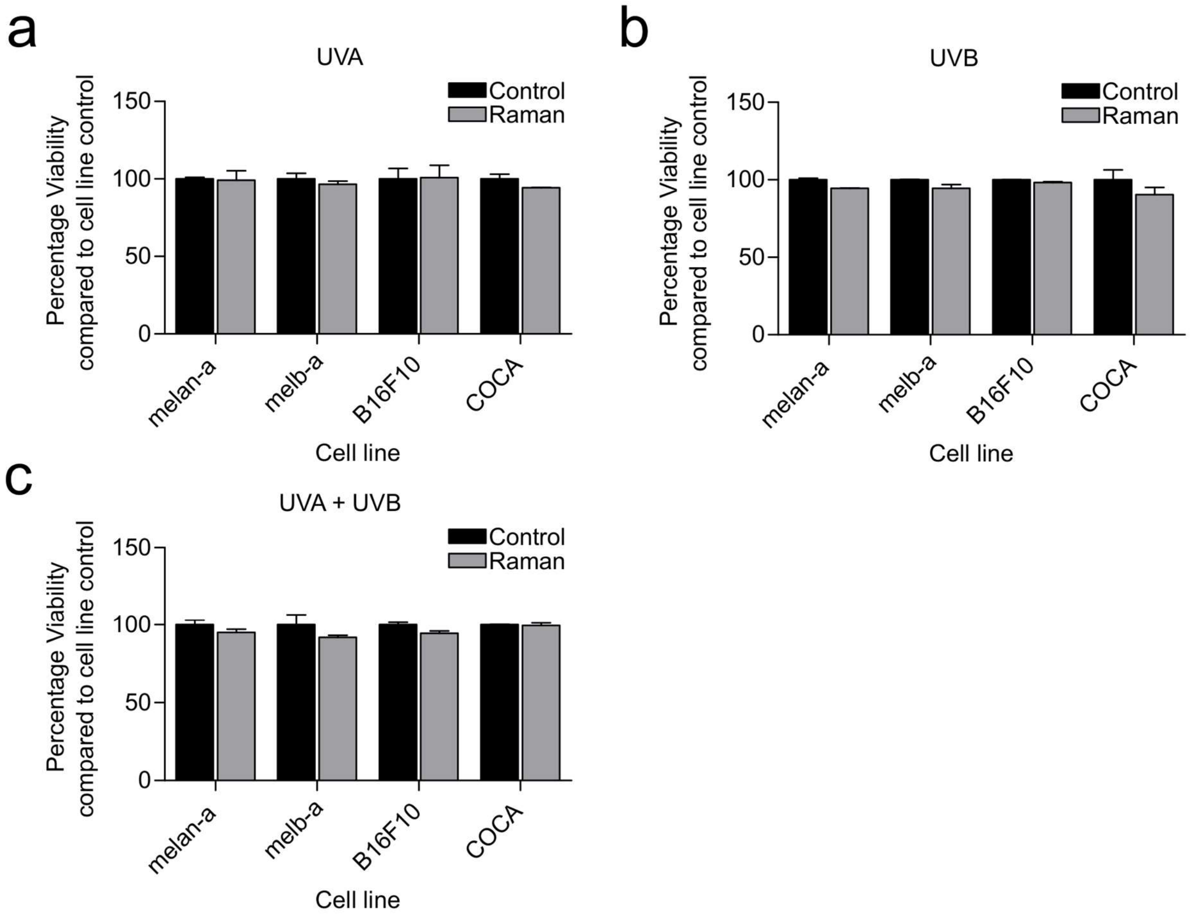


**Supplementary Figure 1. Cell viability post Raman spectral acquisition.** melan-a, melbA, B16F10 and COCA cells were grown on $CaF_2$ disks in duplicate in phenol red free medium for 24 hours before irradiation with 100KJ/m$^2$ UVA, 1000J/m$^2$ UVA and UVB or 100J/m$^2$ UVB before spectra acquired from one disk at 1, 3, 6, 16 and 24 hours post radiation as well as from an untreated control. The other disk was maintained under normal culture conditions as a control. Cell viability was measured using Cell Titer Glo. (A) UVA, (B) UVB and (C) UVA and UVB. Data represented as mean ± 95% CI, N=18.

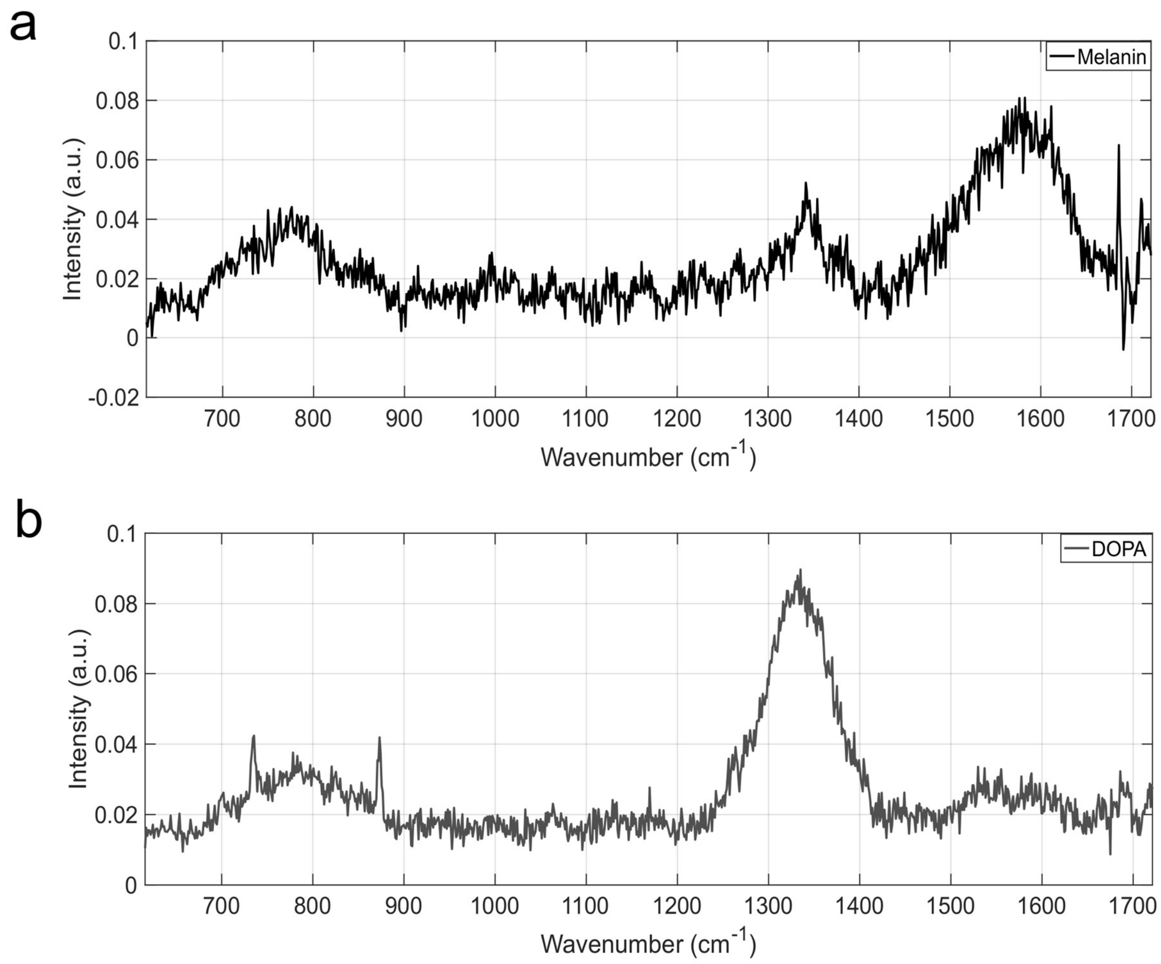


**<u>Supplementary Figure 2. Raman spectra of synthetic melanin and DOPA</u>.** The spectra of melanin and DOPA was acquired using a 785nm laser at a spectral range of 600-1700$cm^{-1}$. Spectra was collected at 2 seconds with 9 accumulations (18 seconds total) at 100% laser power. Data collected was baseline corrected and smoothed using P=0.001, λ=105 before vector normalising.

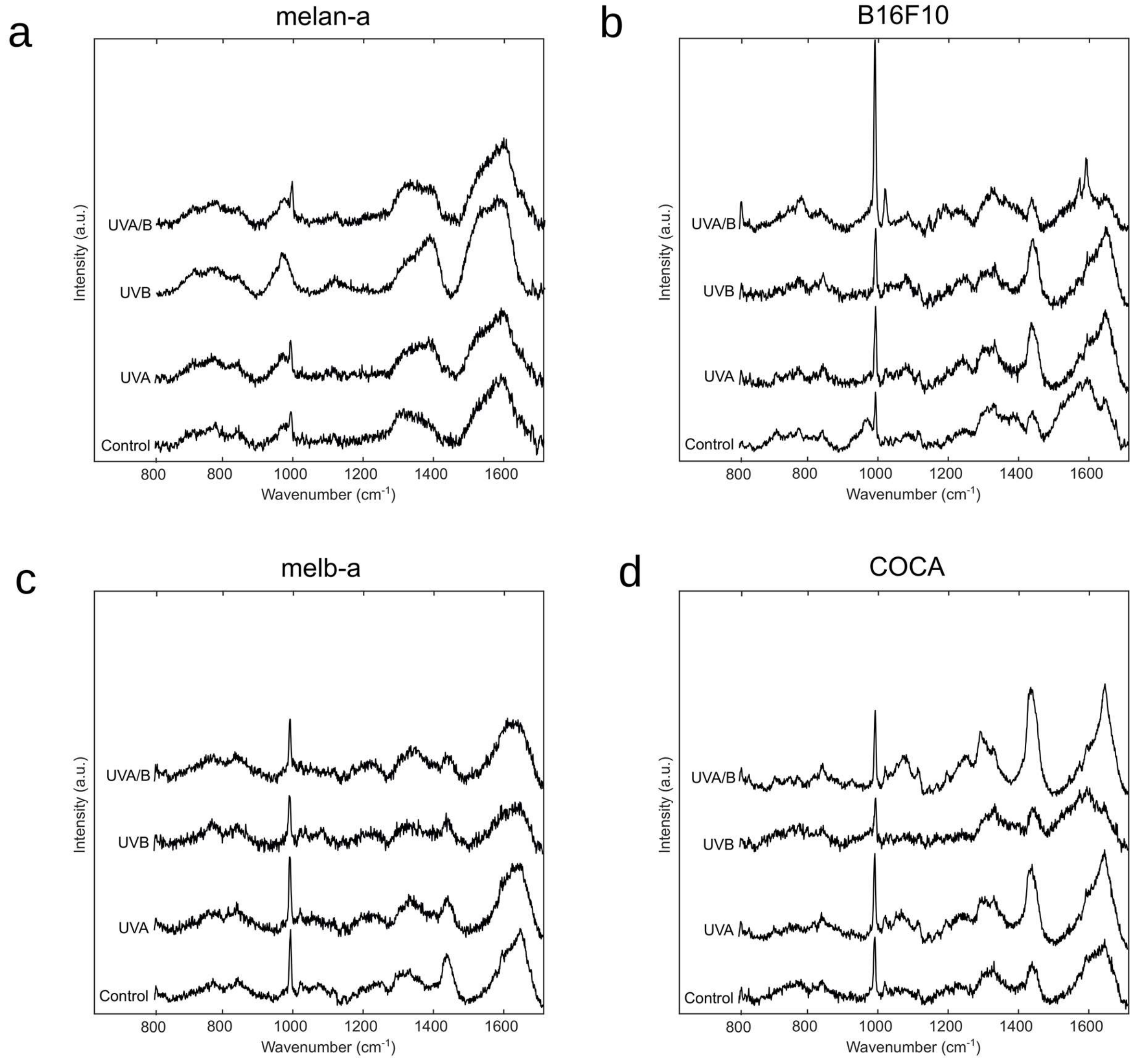


**<u>Supplementary Figure 3. Raman spectra of UVR treated cells.</u>** melan-a, melbA, B16F10 and COCA cells were grown on $CaF_2$ disks and irradiated with 100KJ/m$^2$ UVA, 1000J/m$^2$ UVA and UVB or 100J/m$^2$ UVB before Raman spectra acquired using a 785nm laser at a spectral range of 600-1700cm$^{-1}$, 100% laser power 2 seconds with 9 accumulations (18 seconds total) at 24 hours post irradiation and compare to an untreated control. Data collected was baseline corrected and smoothed using P=0.001, $\lambda$=105 before vector normalising. Raman spectra for each cell line plotted as class mean (n=18) (A) melan-a, (B) B16F10, (C) melbA, (D) COCA.
